# How coupled slow oscillations, spindles and ripples control neuronal processing and communication during human sleep

**DOI:** 10.1101/2023.01.08.523138

**Authors:** Bernhard P. Staresina, Johannes Niediek, Valeri Borger, Rainer Surges, Florian Mormann

## Abstract

Learning and plasticity rely on fine-tuned regulation of neuronal circuits during offline periods. An unresolved puzzle is how the sleeping brain - in the absence of external stimulation or conscious effort – controls neuronal firing rates (FRs) and communication within and across circuits, supporting synaptic and systems consolidation. Using intracranial Electroencephalography (iEEG) combined with multiunit activity (MUA) recordings from the human hippocampus and surrounding medial temporal lobe (MTL) areas, we here show that governed by slow oscillation (SO) up-states, sleep spindles set a timeframe for ripples to occur. This sequential coupling leads to a stepwise increase in (i) neuronal FRs, (ii) short-latency cross-correlations among local neuronal assemblies and (iii) cross-regional MTL interactions. Triggered by SOs and spindles, ripples thus establish optimal conditions for spike-timing dependent plasticity and systems consolidation. These results unveil how the coordinated coupling of specific sleep rhythms orchestrates neuronal processing and communication during human sleep.

## Introduction

How are fleeting experiences transformed into durable memories? Sheltered from external distractors and tasks, sleep constitutes a privileged state for the brain to re-organise and shape neuronal circuits in service of memory formation (Rasch and Born, 2013). Mechanistically, learning and plasticity are governed by two fundamental principles: synaptic consolidation, i.e., long-term potentiation of local circuits afforded by short-latency co-firing (spike-timing dependent plasticity, STDP) and systems consolidation, i.e., the transfer of memory traces across hippocampal-cortical networks via cross-regional communication (Bi and Poo, 2001; Dudai, 2004; Kandel et al., 2014; McGaugh, 2000). An unresolved question is how the sleeping brain regulates neuronal (co-)firing rates to facilitate these forms of consolidation.

Findings from rodent and human electrophysiological recordings point to a potential role of coupled sleep rhythms – slow oscillations (SOs), spindles and ripples - in mediating consolidation processes (Klinzing et al., 2019). SOs reflect fluctuations (< 1 Hz) of membrane potentials, toggling between depolarised up-states and hyperpolarised down-states (Adamantidis et al., 2019; Steriade et al., 1993). Spindles are waxing and waning ∼12-16 Hz oscillations, generated and sustained through thalamocortical interactions (Fernandez and Lüthi, 2020). Ripples are transient high-frequency bursts (∼80-120 Hz in humans), best characterized in the hippocampus but more recently also observed in extrahippocampal/neocortical areas (Buzsáki, 2015; Vaz et al., 2019). Each of these rhythms has been observed in the human medial temporal lobe (MTL) (Clemens et al., 2007; Helfrich et al., 2019; Jiang et al., 2019; Skelin et al., 2021; Staresina et al., 2015) and their interaction has been linked to behavioural expressions of memory consolidation in rodents (Latchoumane et al., 2017; Maingret et al., 2016; Oyanedel et al., 2020), with analogous findings in humans being confined to SO-spindle interactions measured with scalp electroencephalography (EEG) (Helfrich et al., 2018; Muehlroth et al., 2019; Ngo et al., 2013; Schreiner et al., 2021).

Critically, whether and how the interplay of these three sleep rhythms regulates neuronal activity to support synaptic and/or systems consolidation remains unclear. What is the division of labour among SOs, spindles and ripples in controlling local and cross-regional neuronal interactions? In animal models, comprehensive coverage of the MTL and higher-order cortical areas in the same animal/recording session is rare. In humans, data from non-invasive electrophysiological recordings (M/EEG) are restricted to SOs and spindles and localisation to deeper sources remains challenging. Intracranial EEG in epilepsy patients allows recording of SOs, spindles and ripples from the MTL and beyond, but is typically confined to field potentials which are only indirectly related to neuronal firing (Buzsáki et al., 2012). To overcome these limitations, we recorded from the MTL of human epilepsy patients undergoing presurgical monitoring during natural sleep, using depth electrodes furnished with microwires. These microwires capture neuronal firing (multi-unit activity, MUA), allowing us to assess the role of endogenous sleep rhythms in the regulation of local and cross-regional neuronal activity.

## Results

We recorded 20 sessions from 10 participants (range: 1-4 sessions per participant). Depth electrodes were implanted bilaterally, targeting anterior and posterior hippocampus, amygdala, entorhinal cortex and parahippocampal cortex in all participants (Figure 1A). Additional microwires (protruding ∼4 mm from the electrode tips) were used to obtain multi-unit activity (MUA) reflecting neuronal firing. Field potentials capturing slow oscillations (SOs), spindles and ripples were derived from the most medial macro contacts (after bipolar re-referencing). Unless otherwise stated, data were pooled across these MTL contacts (10 per participant) and corresponding microwires (8 per contact). Analyses focused on NREM sleep (stages N2 and N3; see Figure 1B for an example hypnogram and Table S1 for proportions of sleep stages across sessions). SOs, spindles and ripples were detected algorithmically based on previous methods (Ngo et al., 2020). Grand averages of the resulting events are shown in Figure 1C. Note that SO amplitudes are smaller after bipolar re-referencing than after referencing to e.g. linked mastoids, but morphologies and number of detected events are comparable across different re-referencing schemes (Figure S1).

**Figure 1.**
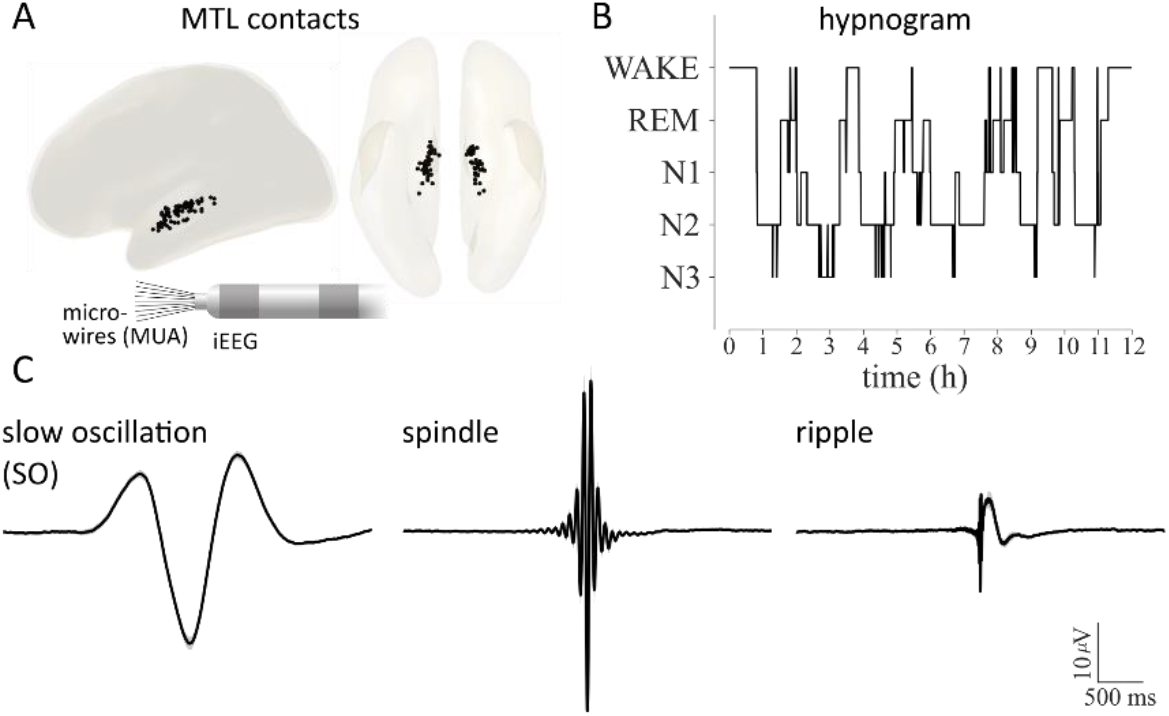
**A**. Locations of iEEG contacts pooled across participants and rendered on a template in MNI space. Inset shows two medial depth electrode contacts (recording the iEEG signal) and the protruding bundle of microwires (recording multiunit activity, MUA). **B**. Example hypnogram from a 12h recording session. Analyses focused on NREM sleep (N2/N3). **C**. Grand average (n=20, ±SEM) of slow oscillations (SOs), spindles and ripples.

### Sequential coupling of SOs, spindles and ripples

To establish the temporal coupling of SOs, spindles and ripples, we first time-locked spindle and ripple centres to SOs, replicating the finding that both event types are nested in SO up-states (Figure 2A, *left*). Importantly, the rate of SO-locked ripples (i.e., a ripple occurring within ±1s of an SO down-state) was significantly higher in the presence of a spindle (same time window) than when no spindle was present (t(19) = 6.47, P < .001; Figure 2A, *right* – statistics conducted across sessions, see Methods for details). To unravel the temporal dynamics between SOs, spindles and ripples in more detail, we repeated the analysis using event onsets instead of event centres. As shown in Figure 2B (*left*), spindle onsets increased in earlier phases of the SO up-state than ripple onsets. Indeed, the maximum event rate (within -2 to 0 s relative to the SO down-state onset) occurred at -451 ms for spindles and at -241 ms for ripples (t(19) = 4.07, P < .001).

**Figure 2.**
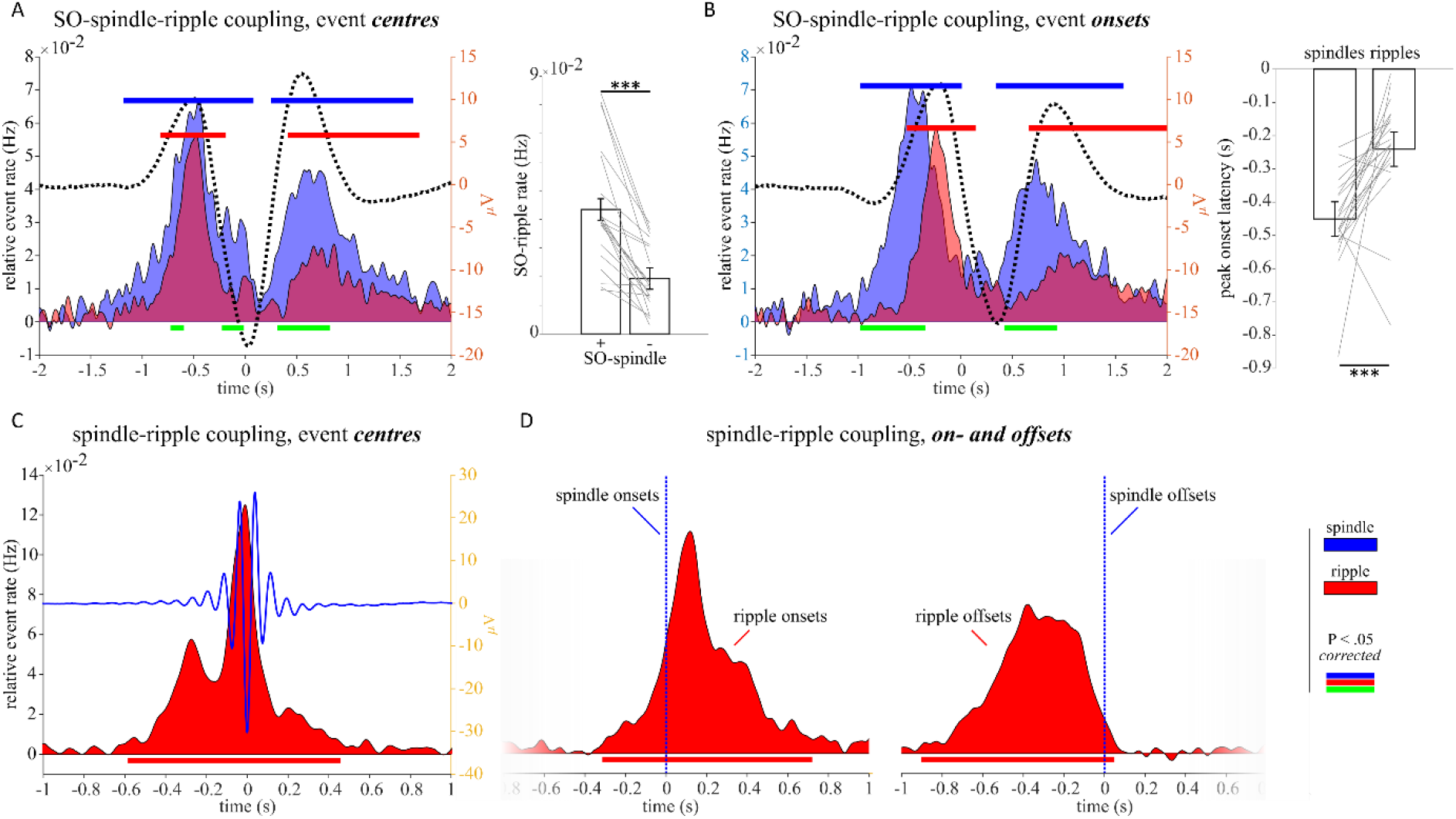
Sequential coupling of SOs, spindles and ripples. **A**. *Left*: Spindle (blue) and ripple (red) rates (centre times) during SOs (dotted black line), relative to a pre-SO baseline period. *Right*: SO-ripple rate (ripple centres within ±1s of SO down-state) as a function of SO-spindle coupling (+: spindle centres within ±1s of SO down-state, -: no spindles within ±1s of SO down-state). Bars show means ± SEM of condition differences. Grey lines reflect individual session data (n=20). **B**. *Left*: Same as **A** but plotting event onsets instead of event centres. *Right*: Latencies (s) of maximal spindle and ripple onset rates from -2s to 0s relative to SO onset. Bars show means ± SEM of condition differences. Grey lines reflect individual session data (n=20). **C**. Ripple rates (centre times, red) during spindles (blue line), relative to a pre-spindle baseline period. **D**. same as **C** but showing ripple onset rates (red) with respect to spindle onsets (vertical blue line; *left*) and ripple offset rates (red) with respect to spindle offsets (vertical blue line; *right*). Horizontal lines: blue … spindle occurrences vs. 0; red … ripple occurrences vs. 0; green … spindle occurrences vs. ripple occurrences, all P < .05 (corrected). *** … P < .001.

To directly test whether spindles might set the timeframes for ripples to occur, we extracted the occurrence of ripple centres, onsets (start times) and offsets (end times) with respect to spindle centres, onsets and offsets. Figure 2C confirms the significant increase of ripple rates around spindle centres, revealing a tendency to occur prior to the spindle centre (i.e., during the ‘waxing’ spindle phase; t(19) = 5.04, P < .001, comparing -1 s to 0 s vs 0 s to 1 s). The on- and offset-locked analysis shown in Figure 2D then corroborated that most ripples coupled to spindles begin after spindle onset and end before spindle offset. For ripple-locked rates of SOs and spindles, see Figure S2. Together, these results show that spindles and ripples cluster in the SO up-state, with spindles increasing the probability and setting the temporal frames for ripples to occur.

### Neuronal firing rates increase across SOs, spindles and ripples

We next turned to the question whether and how neuronal firing rates (FRs) are modulated by SOs, spindles and ripples. Note that previous studies in humans have reported modulation of FRs by SOs (Nir et al., 2011), spindles (Andrillon et al., 2011; Dickey et al., 2021) and (wake) ripples (Tong et al., 2021), but those FRs have not been examined side by side in the same participants, brain regions and behavioural states (e.g., sleep) before. As shown in Figure 3A (single neuron in hippocampus) and 3B (MUA across contacts), all three event types modulated FRs relative to pre-event baseline intervals, but in different manners. During SOs, FRs showed an increase during the up-state and a marked decrease during the down-state, pointing to an active silencing function of SO down-states (FRs below baseline levels). During spindles, FRs increased around the spindle centres and decreased 500 ms before and after the centres, likely reflecting the effect of coupled SO down-states (see Figures 2A and S3). Finally, FRs showed a pronounced increase during ripples, exhibiting a ∼500 ms ramp-up period prior to the ripple start. To quantify the stepwise increase in FRs across SOs, spindles and ripples, we derived – for each session - the maximum FRs within ± 2 s of the three event centres. As illustrated in Figure 3C, there was a significant increase in max FRs from SOs to spindles (t(19) = 2.21, P = .040) and from spindles to ripples (t(19) = 3.96, P < .001).

**Figure 3.**
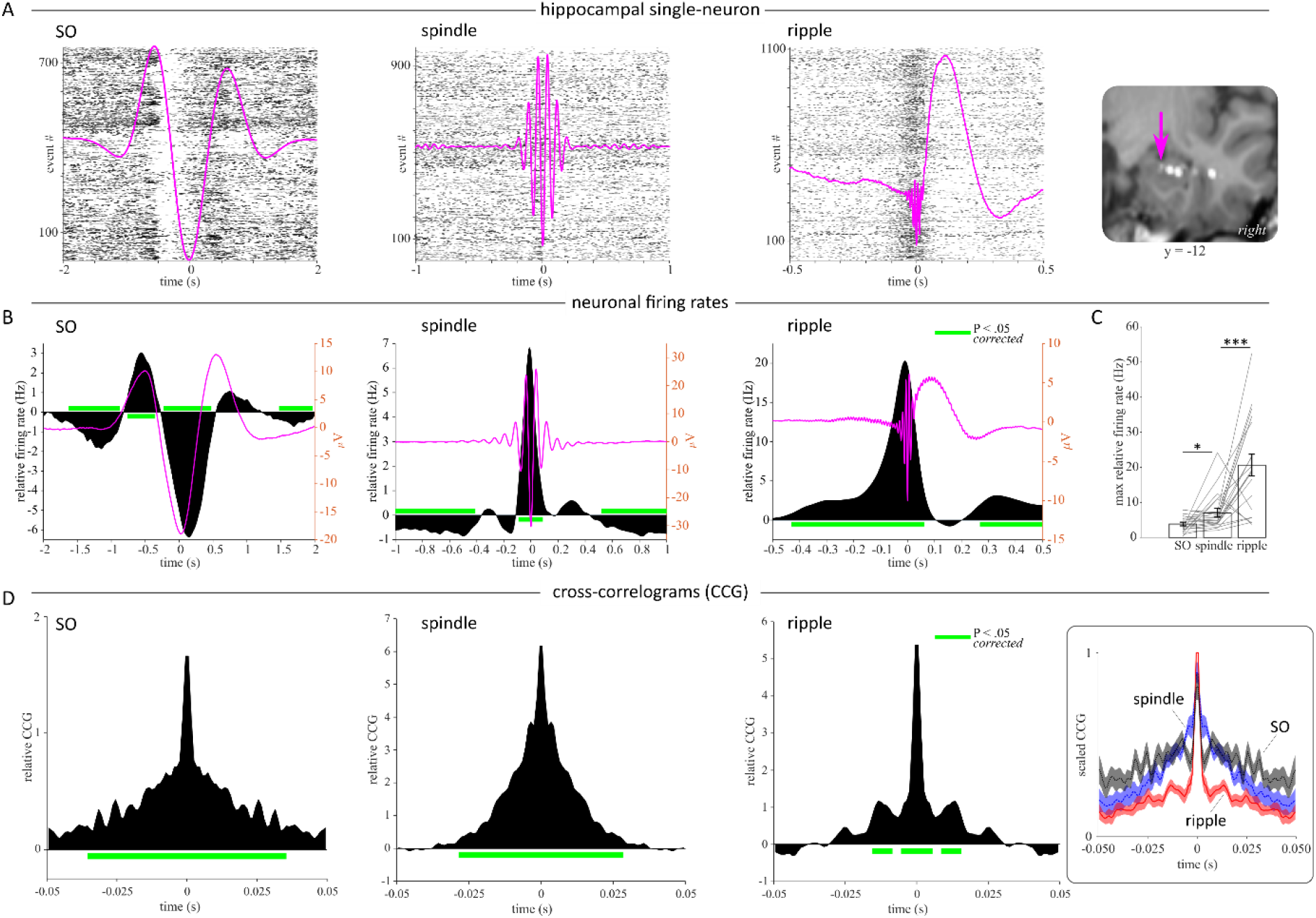
Modulation of neuronal (co-)firing rates (FRs) by SOs, spindles and ripples. **A**. Single participant/single neuron example. Raster plots show action potentials across time (x axis) for individual events (y axis) and session-specific SOs, spindles and ripples (magenta). ERPs were bandpass filtered in the SO and spindle detection range, respectively, and from .1 Hz - 120 Hz for ripples to preserve the sharp-wave component for visualisation. *Right*: MRI-CT scan illustrating the macro contact and the microwire bundle (arrow) from which this single unit was isolated. Y-coordinate refers to MNI space. **B**. FRs relative to pre-event baselines (black) and grand-average ERPs (magenta) for SOs (*left*), spindles (*middle*) and ripples (*right*). **C**. Maximal FRs per session (n=20, grey lines), illustrating the stepwise increase across SOs, spindles and ripples. Bars show means ± SEM of conditions. **D**. Event-locked cross-correlograms (CCGs) of neuronal firing among pairs of microwires, relative to ‘shift-predictors’ and non-event surrogates. *Right*: CCGs for SOs (black), spindles (blue) and ripples (red) scaled between 0 and 1, illustrating the stepwise narrowing of co-firing windows. Horizontal green line … FRs vs. 0, P < .05 (*corrected*). * … P < .05, *** … P < .001.

### Neuronal co-firing during SOs, spindles and ripples

As mentioned above, one central mechanism driving learning-related changes in cell assemblies is short-latency co-firing, capable of inducing long-term potentiation (LTP) via spike-timing dependent plasticity (STDP) (Bi and Poo, 1998). A recent report showed that spindles in lateral temporal cortex group co-firing within 25 ms (Dickey et al., 2021), but how this effect relates to potential co-firing during SOs and ripples is unclear. To examine co-firing patterns during SOs, spindles and ripples, we derived cross-correlograms (CCGs) in ±50 ms windows centred on event maxima. CCGs were calculated for all pairwise combinations of microwires on a given bundle (resulting in symmetrical CCGs), including only wires that showed a minimum FR of 1 Hz during all NREM sleep. Resulting CCGs were corrected in two steps. First, we subtracted ‘shift-predictor’ CCGs, reflecting the cross-correlation of wire 1 FR during event n with wire 2 FR during event n+1, thereby accounting for overall FRs during a particular event type (Steinmetz et al., 2000). Second, we subtracted CCGs derived from matched non-event surrogates (also shift-predictor-corrected). In the resulting CCGs, values above zero thus signify highly event-specific co-firing in local assemblies. As shown in Figure 3D, all three event types elicited significant neuronal co-firing. Importantly though, the temporal windows of co-firing displayed a stepwise decrease across SOs, spindles and ripples. Significant co-firing spanned a range of 35 ms for SOs, 28 ms for spindles and 5 ms for ripples, with a second peak between ∼9 and 15 ms (reflecting a ∼70-110 Hz oscillatory cycle). The stepwise narrowing of CCGs is further highlighted in Figure 3D (*right*), where all three CCGs were scaled between 0 and 1 within session. Together, these findings suggest that ripples are most apt in creating conditions conducive to STDP.

### Relationship between firing rates and event occurrences across SOs, spindles and ripples

The strong increase in FRs during ripples (Figure 3B) raises the question whether the SO- and spindle-related FRs might merely reflect ripple-related FRs, given that ripples are coupled to SOs and spindles (Figure 2A). Conversely, genuine FR increases during SOs and spindles might trigger ripple occurrences by mediating the observed ramp-up preceding ripples (starting ∼500 ms prior to ripple centres; Figure 3B). To adjudicate between these two scenarios, we first conducted the event-locked FR analysis again, but separating event types of interest (e.g., spindles, ‘seed’) based on the presence or absence of another event type (e.g., ripples, ‘target’). Presence/absence was coarsely defined as algorithmically detected target event centres occurring within ±1 s of the seed event centre. As shown in Figure 4A, both SOs and spindles showed enhanced FRs when ripples were present (and vice versa, Figure S3). Importantly though, FRs in the SO up-states and around spindle centres also exhibited significant increases when no ripples were present (for all pairwise seed/target combinations, see Figure S3). This result indicates that ripples are not the (sole) driver of FR increases during SOs and spindles.

**Figure 4.**
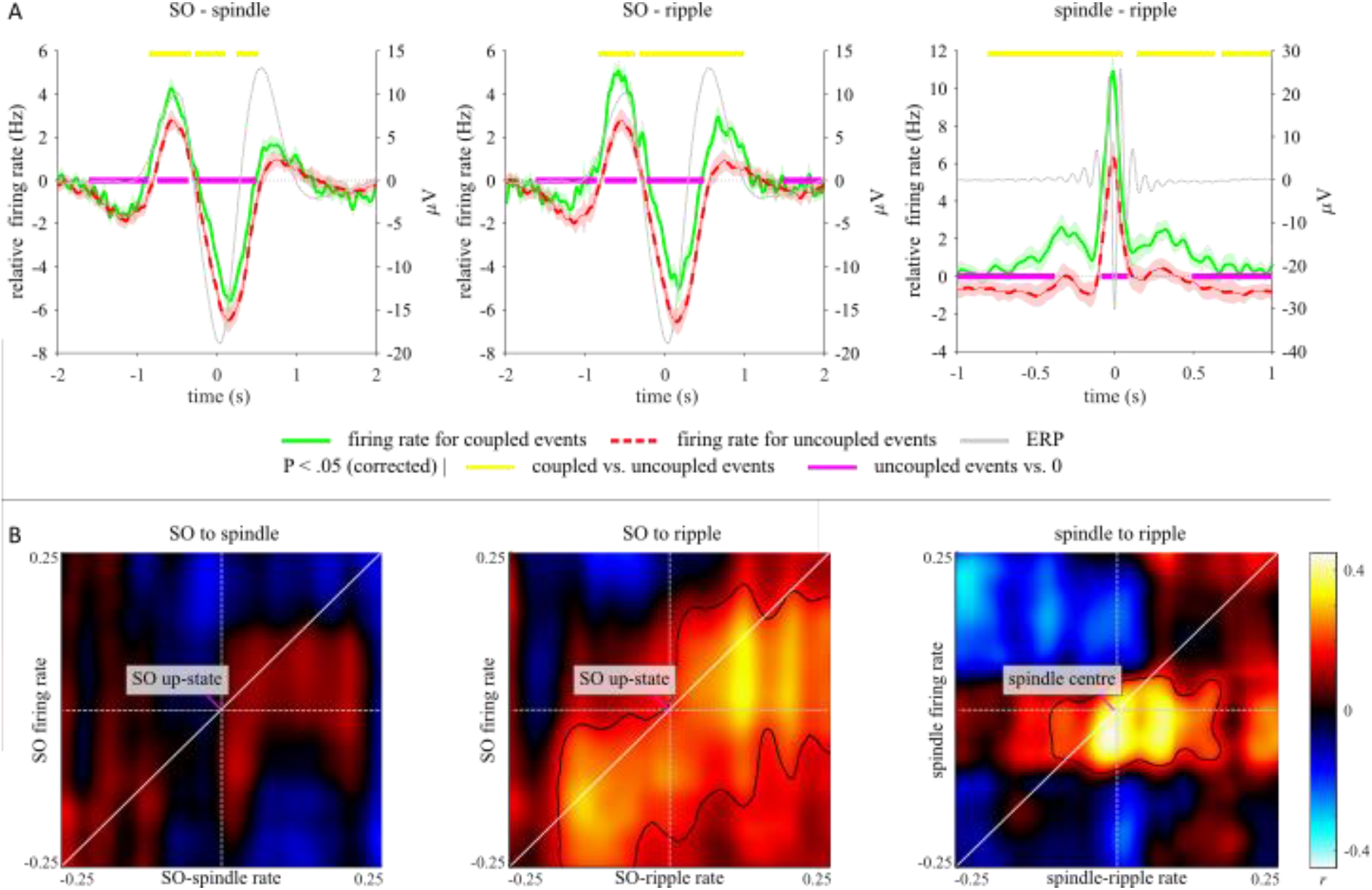
Relationship between neuronal firing rates (FRs) and event occurrences. **A**. FRs as a function of event contingencies (target event centre within ±1 s of seed event centre). Green/red lines represent mean baseline-corrected FR across sessions (n=20) ± SEM of condition differences (coupled vs. uncoupled events). Grey lines represent grand average ERPs of seed events (irrespective of target presence). Note that FRs are elevated for all coupled events, but SOs and spindles without ripples also elicit significant above-baseline FRs. **B**. Time-by-time correlations between seed event FR and target event occurrences. Correlations (Pearson) are shown for 500 ms intervals between SO FRs and SO-locked spindle rate (*left*), SO FRs and SO-locked ripple rate (*middle*) and spindle FR and spindle-locked ripple rate (*right*). Below diagonal: firing rates precede event occurrences, above diagonal: firing rates follow event occurrences. Contour … P < .05 (corrected).

To address the second question, i.e., whether SO- and spindle-related FRs might predict ripple occurrences, we computed – separately for each session – a time by time correlation between (i) seed event FRs and (ii) target event rates across the 10 MTL contacts of each participant. In other words, does a contact that shows greater FRs during spindles also show greater spindle-locked ripple rates? The resulting correlation maps (n=20) were then tested against 0 via non-parametric cluster permutation tests. As shown in Figure 4B, we observed no association between FRs during SO up-states and SO-locked spindle rates. However, FRs during SO up-states were positively correlated with SO-locked ripple rates. This relationship was even more pronounced for FRs around spindle centres and spindle-locked ripples rates. Critically, significant correlations were seen below the diagonal, i.e., earlier SO/spindle FRs predicted later ripple rates. Together, these results suggest that ripples are (at least in part) triggered by gradual increases in FR afforded by SOs and spindles.

### MTL network dynamics linked to SOs, spindles and ripples

Lastly, systems consolidation relies on inter-regional information transfer beyond local cell assemblies (Alvarez and Squire, 1994; Frankland and Bontempi, 2005). We thus set out to examine the role of SOs, spindles and ripples in cross-regional interactions among the separate MTL regions targeted in our recordings (anterior hippocampus, posterior hippocampus, amygdala, entorhinal cortex and parahippocampal cortex, Figure 5A). Note that bipolar referencing was employed for this dataset, thereby mitigating the risk of spurious effects reflecting volume conduction.

**Figure 5.**
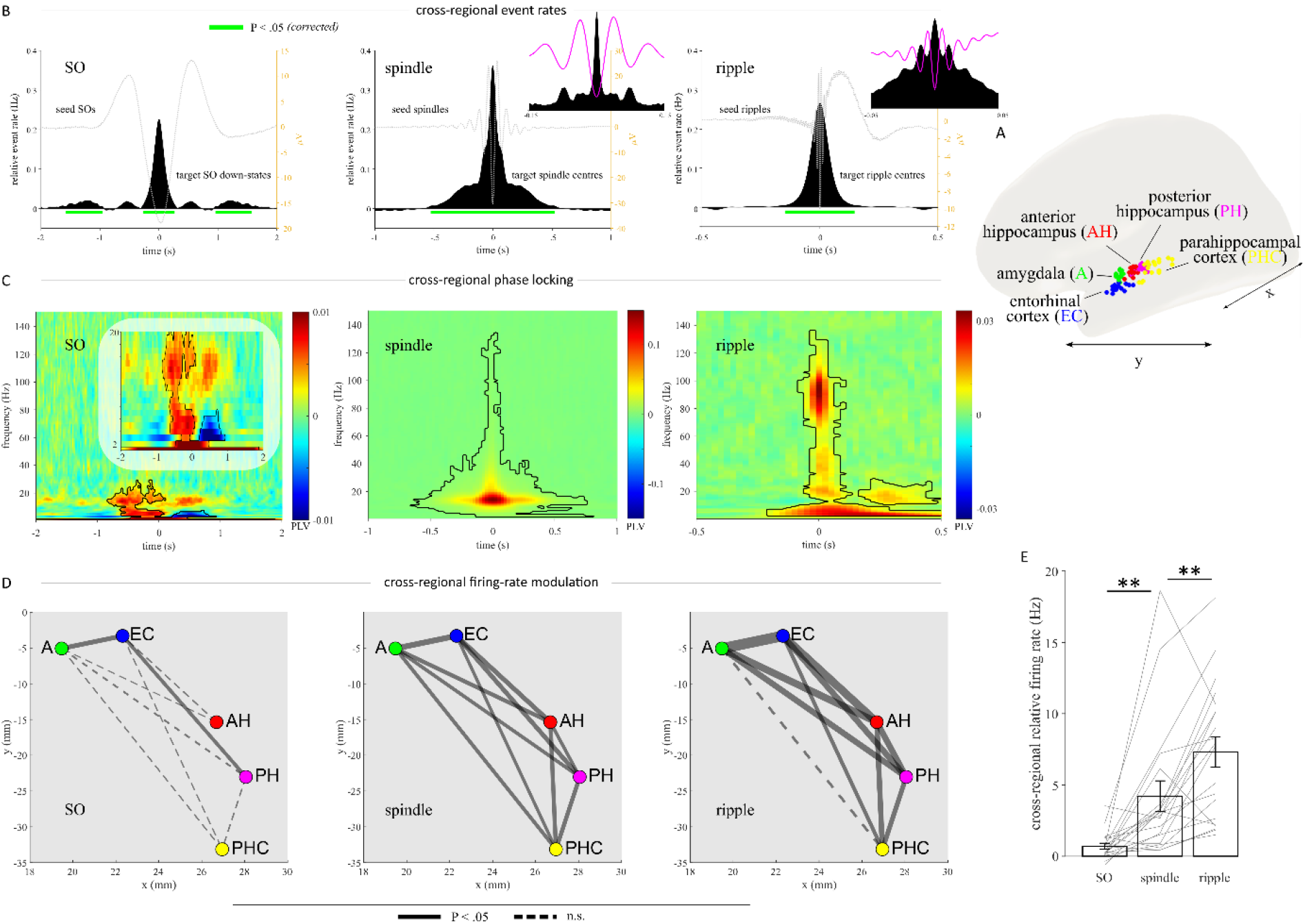
Network dynamics. **A**. Locations of anterior hippocampus (AH, red), posterior hippocampus (PH, magenta), amygdala (A, green), entorhinal cortex (EC, blue) and parahippocampal cortex (PHC, yellow). MNI coordinates were averaged across participants and projected onto the left hemisphere of a standard template. **B**. Cross-regional event rates relative to a pre-event baseline window, showing significant co-occurrences of SOs, spindles and ripples. Insets show data after applying 10 ms instead of 100 ms smoothing, highlighting the temporal precision of co-occurrences. **C**. Time-frequency-resolved phase locking values (PLV) between a seed region and the four same-hemisphere target regions, relative to a pre-event baseline and averaged across sessions (n=20). Contour: P < .05 (corrected). **D**. Cross-regional event-locked firing rates (FRs) for SOs, spindles and ripples (averaged across a 100 ms window centred on the event maximum). Nodes reflect projections of MTL coordinates (MNI space) on the XY plane. Edge widths reflect effect sizes (t values) of bidirectional increases in FR relative to a pre-event baseline. Solid line: significant bidirectional up-regulation of FRs. Dashed line: not significant. **E**: All edges collapsed. Bars show mean ± SEM of conditions. ** … P < .01.

In a first step, we assessed the extent of cross-regional event occurrences, i.e., the likelihood of e.g. a ripple also occurring in region B (target region) when a ripple is detected in region A (seed region). All combinations of same-hemisphere contact pairs were collapsed (resulting in symmetrical histograms), and target event rates were again compared to a pre-event baseline interval. As illustrated in Figure 5B, all three event types were coupled across MTL regions. It is worth highlighting the precise cross-regional locking to the seed events for spindles and ripples (*insets*), indicating that the majority of other spindle/ripple centres occurred either in the same or the next cycle, thereby optimising conditions for mutual communication (Fell and Axmacher, 2011).

To corroborate and expand on the finding of cross-regional communication, we calculated pairwise phase-locking values (PLVs), reflecting consistent phase differences of two regions across events (Lachaux et al., 1999). This was done in a time- and frequency-resolved manner, centred around SOs, spindles and ripples, and statistically compared to pre-event baselines (again collapsing across same-hemisphere contact pairs). As shown in Figure 5C, this analysis confirmed that all three event types elicited significant PLV in the seed event’s frequency range (see previous analysis). It is interesting to note that regions also coupled in the spindle band during SOs and following ripples, extending a recent finding that spindles mediate hippocampal-cortical communication during ripples (Ngo et al., 2020).

As a last step, we assessed the extent to which the observed functional connectivity translates to cross-regional modulation of neuronal firing rates, e.g., a ripple in region A leading to an increase in FR in region B. Specifically, we derived – for each pair of same-hemisphere MTL contacts – the baseline-corrected FRs in the target region (averaged across a 100 ms window centred on the seed region’s event maximum). Resulting connection strengths across the MTL (t values combined across both seed/target directions for a given pair and across both hemispheres) are shown in Figure 5D. Collapsing all connection nodes illustrates the stepwise increase in cross-regional FR modulation across SOs, spindles and ripples (Figure 5E). Taken together, these results reveal that SOs, spindles and ripples are functionally coupled across MTL regions. This coupling leads to fine-tuned network modulation of neuronal firing rates (most strongly during ripples), putatively well-suited to support systems-level consolidation.

## Discussion

Our findings elucidate the mechanisms through which slow oscillations (SOs), spindles and ripples control neuronal firing rates (FRs) and communication during sleep, establishing conditions conducive to synaptic and systems consolidation. While recent work has converged on the importance of these sleep rhythms’ (co-)occurrence for memory consolidation (Diekelmann and Born, 2010; Klinzing et al., 2019), their division of labour in the process has remained elusive. Our data suggest that SO up-states first establish a coarse time window for spindles and ripples to coincide, consistent with previous reports of triple-nesting of these events (Clemens et al., 2007; Helfrich et al., 2019; Jiang et al., 2019; Latchoumane et al., 2017; Maingret et al., 2016; Oyanedel et al., 2020; Staresina et al., 2015). Importantly, spindles enhance the likelihood (Figure 2A) and set a more fine-grained temporal frame for ripples to occur (Figure 2C), akin to a relay function of spindles between SOs and ripples. Mechanistically, our data suggest that SO up-states and – to a greater extent – the waxing phase of spindles elevate FRs to a threshold at which ripples are triggered (Figure 4B). This results in an exponential increase in FRs (Figure 3B and C) and concomitant synchronisation in (i) local cell assemblies (Figure 3D) and (ii) across the MTL (Figure 5).

The enhancement of neuronal FRs and ripple rates when SOs and spindles are coupled (Figures 2A, 4A, S2) dovetails with a series of recent findings linking SO-spindle coupling to physiological and behavioural manifestations of memory consolidation (Helfrich et al., 2018; Muehlroth et al., 2019; Niethard et al., 2018; Schreiner et al., 2021; Silversmith et al., 2020). Moreover, optogenetic enhancement of spindles in mice was found to elicit hippocampal ripples and memory improvements, particularly when stimulation occurred during SO up-states (Latchoumane et al., 2017). That said, the facilitation of ripples is unlikely to be the sole function of SOs and spindles. Likewise, SOs and spindles are clearly not the only means through which ripples can be triggered – for one, ripples are readily observed during waking states (Chen et al., 2021; Jadhav et al., 2012; Vaz et al., 2019). However, in the absence of external input and conscious control, coupled SO-spindle events constitute a controlled yet effective mechanism of gradually elevating neuronal FRs and thereby triggering ripples (Csicsvari et al., 2000; Sirota et al., 2003).

Apart from grouping spindles and ripples in their up-states, one striking feature of the SO-locked analysis was the active inhibition of FRs (below baseline levels) during down-states (Figure 3B). This effect (also referred to as OFF-periods) is well documented across species (Cash et al., 2009; Nir et al., 2011; Steriade et al., 1993; Vyazovskiy and Harris, 2013) and points to a dynamic alternation between active consolidation processes during up-states and homeostatic recalibration and/or pruning of irrelevant circuits during down-states (Tononi and Cirelli, 2006; Vyazovskiy et al., 2009).

Spindle-ripple coupling has been established in animal and human intracranial recordings (Clemens et al., 2007; Dickey et al., 2022; Jiang et al., 2019; Ngo et al., 2020; Siapas and Wilson, 1998; Sirota et al., 2003; Staresina et al., 2015). A consistent finding in these (and our current) data is that spindles nest ripples after their onset and just prior to their maximum (Clemens et al., 2011; Oyanedel et al., 2020; Staresina et al., 2015). In light of the ability of spindles to synchronise wide-ranging neuronal networks (Andrade et al., 2011; Contreras et al., 1997; Fernandez and Lüthi, 2020), an intriguing possibility is that spindles not only drive ripple emergence during their waxing phase, but that the ongoing synchronisation during the waning phase supports the inter-regional transfer of information reactivated during ripples (Diekelmann and Born, 2010). Indeed, we observed increased spindle band phase-locking after ripples across the MTL (Figure 5C), extending our recent finding of post-ripple coupling between the hippocampus and scalp recordings (Ngo et al., 2020). Although evidence for ripple-mediated memory reactivation during sleep is still lacking in humans, it has been firmly established in rodent work (Buzsáki, 2015; Diba and Buzsáki, 2007; Dupret et al., 2010; Girardeau et al., 2009; Lee and Wilson, 2002).

Memory consolidation ultimately reflects adaptive changes in brain structure and function (Dudai, 2004; McGaugh, 2000). On a synaptic level, such changes can be afforded by long-term potentiation and spike-timing-dependent plasticity (STDP), elicited by short-latency co-firing of participating neurons (Caporale and Dan, 2008; Markram et al., 1997; Whitlock et al., 2006). Ripples reflect a surge in local circuit synchronization, ideally poised to induce such synaptic changes (Buzsáki, 2015; Girardeau and Zugaro, 2011). Our results reveal that relative to SOs and spindles, ripples indeed create the narrowest (<10 ms) time windows of neuronal co-firing (Figure 3D), thus supporting the notion that ripples are a viable mechanism to induce synaptic consolidation in humans.

Finally, the persistence of memories is thought to rely on their distribution across hippocampal-cortical networks (‘systems consolidation’) (Alvarez and Squire, 1994; Buzsáki, 1996; Diekelmann and Born, 2010; Dudai, 2004; Frankland and Bontempi, 2005; Roüast and Schönauer, 2022). In rodents and non-human primates, ripples have been shown to impact activation in and connectivity among long-range cortical and subcortical brain networks (Logothetis et al., 2012; Nitzan et al., 2022). While we observed that SOs, spindles and ripples are all synchronised across the MTL (Figure 5A and B), cross-regional modulation of FRs was again strongest during ripples (Figure 5C). Together, these results suggest that whereas SOs and particularly spindles open and maintain channels for cross-regional communication, ripples provide further means to effectively forge local and brain-wide functional networks.

While our results are consistent with a role of SO-spindle-ripple coupling in promoting synaptic and systems consolidation, a clear shortcoming of the current study is the lack of a proper assay to capture behavioural expressions of memory consolidation. It is a daunting challenge in patient work to devise such an assay, as robust conclusions ideally require not only large sample sizes to allow cross-participant correlations, but also multiple sessions within a participant to compare nights of ‘better’ consolidation with nights of ‘worse’ consolidation. Another caveat is that the raw numbers of event (co-)occurrences as derived here are somewhat arbitrary, as they heavily rely on the parameters set in the detection algorithms. For instance, we have shown previously (Staresina et al., 2015) that e.g. the proportion of SO-triggered spindles which also contain ripples increases from 6 to 10% when minimally relaxing the detection thresholds (setting the ripple detection threshold from the top 1% to the top 2% amplitude). That said, examination of relative event rates after normalisation to a pre-event baseline or to matched surrogate events is still highly informative, as the detection parameters (and ensuing miss/false alarm rates) are held constant across target and baseline/control epochs. By the same token, the ‘absence’ of a co-occurring event (Figure 4A, Figure S3) warrants interpretive caution, as subthreshold events might have been missed by the detection procedure.

To conclude, we show that SOs, spindles and ripples systematically interact to control neuronal firing rates and communication during NREM sleep. Ignited by SO up-states, spindles set the temporal frames for ripples to occur. In turn, ripples lead to a surge in neuronal firing and drive short-latency coactivation in local assemblies, fostering conditions permissive of STDP/LTP. Finally, sleep rhythms are synchronised across the medial temporal lobe, facilitating cross-regional neuronal communication thought to underly systems consolidation.

## Methods

### Participants and recordings

Macro- and simultaneous microwire recordings were performed in 20 sessions from 10 participants (range: 1-4 sessions per participant) undergoing invasive presurgical seizure monitoring for treatment of medically refractory epilepsy (5 male, 5 female, all right-handed, mean age = 39.9 years, range = 20-62 years). The study conformed to the guidelines of the Medical Institutional Review Board at the University of Bonn, and participants provided written informed consent. Depth electrodes were implanted bilaterally, targeting anterior and posterior hippocampus (hippocampal head and body, respectively), amygdala, entorhinal cortex and parahippocampal cortex in all participants (Figure 1A). Depth electrodes were furnished with bundles of nine microwires each (eight high-impedance recording electrodes, one low-impedance reference, AdTech, Racine, WI) protruding ∼4 mm from the electrode tips. The differential signal from the microwires was amplified using a Neuralynx ATLAS system (Bozeman, MT), filtered between 0.1 Hz and 9000 Hz, and sampled at 32 kHz.

Multiunit activity (MUA) reflecting neuronal firing was obtained from these microwires using the Combinato package (Niediek et al., 2016). In brief, spikes were identified from each wire independently via a thresholding procedure and extracted after bandpass filtering (300 to 3000 Hz). Combinato’s default procedure for artefact removal was applied: 500 ms time bins containing more than 100 events were excluded; events exceeding an amplitude of 1 mV were removed; 3 ms time bins with coinciding events on more than 50% of all channels were excluded. Remaining spikes were spike-sorted, and artefact clusters were identified. Combinato’s default parameters were used in each step. All non-artefact clusters were then merged on each channel separately. All FR analyses reported here were derived from these MUA signals.

Field potentials capturing slow oscillations (SOs), spindles and ripples were derived from the proximal (most medial) macro contacts, thus mitigating contamination of the intracranial EEG signal by high-frequency action potentials (Liu et al., 2022). Unless otherwise stated, data were pooled across all MTL regions and microwires. Each macro contact was re-referenced in a bipolar fashion, subtracting the signal from the neighbouring contact on the same electrode. This procedure was chosen to safeguard against signal spread from adjacent MTL areas, which would be a particular concern for connectivity analyses. The effect of different referencing schemes (bipolar, white matter contact, linked mastoids) on the resulting event numbers and morphologies is shown in Figure S1. Event densities separated by MTL region are shown in Figure S4.

For polysomnography (PSG), additional surface electrodes were applied according to the 10-20 system alongside EOG and EMG electrodes. Sleep staging was performed according to AASM guidelines (Iber, 2007) and all analyses were confined to NREM sleep (N2 and N3). Figure 1B shows an example hypnogram, and proportions of sleep stages across sessions are listed in Table S1.

### Artefact rejection and event detection

Data were analysed in MATLAB (version R2022a) using FieldTrip functions (version 20201229) (Oostenveld et al., 2011). Prior to sleep event detection, each contact was subjected to preprocessing and artefact detection. Preprocessing included down-sampling to 1 kHz, removing 50 Hz line noise and harmonics up to 200 Hz via +/- 1 Hz bandstop filters, and 0.1 Hz high pass filtering to remove slow signal drifts. The iEEG signal was inverted so that positive peaks reflect up-states. For artefact detection, two copies of the raw signal were created – one after applying a 250 Hz high-pass filter and one after taking the first derivative of the data (reflecting signal gradients). The three signals (raw, hp-filtered, gradient) were z-scored within each sleep stage, and a data point was classified as artefactual if it exceeded a z score of 6 in any one of the three signals or a zscore of 4 in the raw signal as well as in any of the two other signals (hp-filtered, gradient). Artefacts less than 3 s apart were merged, and artefactual samples was additionally padded by 1 s on each side.

SOs, spindles and ripples were detected algorithmically based on previous methods (Ngo et al., 2020; Staresina et al., 2015). In brief, for SO detection, the signal was first bandpass-filtered between .3 and 1.25 Hz. Second, all zero-crossings were determined in the filtered signal, and event duration was determined for SO candidates as time between two successive positive-to-negative zero-crossings (i.e., a down-state followed by an up-state). Events that lasted between 0.8 and 2 s entered the next step. Third, event amplitudes were determined for the remaining SO candidates (trough and trough-to-peak amplitude). Events in which both amplitudes exceeded the mean plus 1 standard deviation of all candidate events were considered SOs. SO-locked analyses were either based on the event onset (position of positive-to-negative zero crossing/up-to down-state transition), the event centre (maximal trough after the onset, i.e., down-state) or the event maximum (maximal peak after the onset, i.e., up-state).

For spindle detection, the signal was band-pass filtered at 12–16 Hz, and the root mean square (RMS) signal was calculated based on a 200 ms window followed by an additional smoothing with the same window length. A spindle event was identified whenever the smoothed RMS signal exceed a threshold, defined by the mean plus 1 standard deviation of the RMS signal across all NREM data points, for at least 0.4 s but not longer than 3 s. Time points exceeding an upper threshold determined by the mean RMS signal plus 9 times its the standard deviation were excluded. Lastly, spindles were required to exhibit a minimum of 6 cycles in the raw EEG signal. The onset (start) and offset (end) of spindles were defined as the upward and downward threshold crossings, respectively. Unless otherwise noted, spindle centres were defined as the maximal trough.

Detection of ripples followed the same procedure, except that the iEEG signal was bandpass-filtered from 80 to 120 Hz, and both RMS calculation and smoothing were based on 20 ms windows. Detection and upper cut-off threshold were defined by the mean of the RMS-signal plus 3 or 9 times the standard deviation, respectively. Potential ripple events with a duration shorter than 38 ms (corresponding to 3 cycles at 80 Hz) or longer than 200 ms were rejected. Additionally, all ripple events were required to exhibit a minimum of three cycles in the raw EEG signal. Unless otherwise noted, ripple centres were defined as the maximal trough.

### Anatomy

Post-implantation MRI-CT scans were available for 9 of the 10 participants. Scans were normalised to MNI space using SPM12. Contact locations were then manually identified to create group-level representations of target locations (Figure 1A). To visualise the MTL network (Figure 5), XYZ coordinates were averaged across participants.

### Analyses and statistics

Analyses of event occurrences and firing rates were based on peri-event time histograms (PETHs), with 1 ms bin sizes. Event/firing rates were converted to Hz and temporally smoothed with a 100 ms Gaussian kernel unless otherwise noted. Resulting histograms were then corrected to pre-event baseline intervals (−2.5 s to -2 s for SOs and spindles, -1.5 s to -1 s for ripples). For the CCG analysis, additional non-event surrogates were derived. For instance, for each participant’s n observed ripple events, we derived n non-ripple events, i.e., artefact-free NREM epochs matching the duration of each individual event including an additional padding of 1.5 s before and after in which our ripple detection algorithm did not indicate the presence of a ripple (irrespective of the presence/absence of spindles or SOs). Furthermore, to ensure that signal properties were maximally matched between target events and surrogates, control events were only drawn from a 10 min time window before and after the corresponding ripple event. The probability underlying the randomized selection of control events within such a 10 min interval was modulated according to a normal distribution. Epochs once assigned to control events were discarded from subsequent iterations to exclude overlapping non-events.

Analyses were performed separately for each MTL contact, pooling firing rates across the 8 microwires in case of MUA analyses (except for the CCG analysis shown in Figure 3D). Session-specific results were obtained by averaging results across contacts (except for cross-contact correlations shown in Figure 4C and connectivity analyses shown in Figure 5), weighing each contact by the number of seed events of that contact. Final statistical analyses were performed across sessions (n=20).

For the CCG analysis shown in Figure 3D, we used FieldTrip’s ft_spike_xcorr function. Time windows to derive CCGs were equated across events and set to 150 ms centred on event maxima (to capture the SO up-state), including positive and negative lags up to 50 ms. Bin size was 1 ms, and resulting CCGs were smoothed with a 5 ms Gaussian kernel.

For the cross-regional phase-locking-value (PLV) analysis shown in Figure 5, time-frequency representations (TFRs) were extracted centred on target event maxima (using FieldTrip’s mtmconvol function) for frequencies from 1 Hz to 150 Hz in steps of 1 Hz, using a sliding Hanning-tapered window advancing in 25 ms steps. The window length was frequency-dependent, such that it always comprised a full number of cycles but at least five cycles and at least 100 ms, ensuring reliable phase estimates for higher frequencies for which five cycles would result in too short windows.

To correct statistical analyses for multiple comparisons, a cluster-based permutation procedure was applied as implemented in FieldTrip, using 1,000 permutations, a cluster threshold of P < .05 and a final threshold for significance of P < .05 (all two-tailed).

## Acknowledgments

This work was supported by an ERC Consolidator grant (101001121) to B.P.S and grants from the DFG (MO 930/4-2, SFB 1089, SPP 2205), BMBF (031L0197B) and a NRW Network Grant (iBehave) to F.M.

## Author contributions

Conceptualization, B.P.S. and F.M.; Methodology, B.P.S., J.N.; Software, B.P.S. and J.N.; Formal Analysis, B.P.S.; Investigation, J.N.; Writing – Original Draft, B.P.S.; Writing – Review & Editing, J.N. and F.M.; Funding Acquisition, B.P.S. and F.M.; Surgical Procedures; V.B. and F.M.; Patient Recruitment; R.S.

## Data availability

Data and analysis scripts to reproduce the main results will be shared on the Open Science Framework (OSF) upon publication.

## Supplemental results

**Figure S1.**
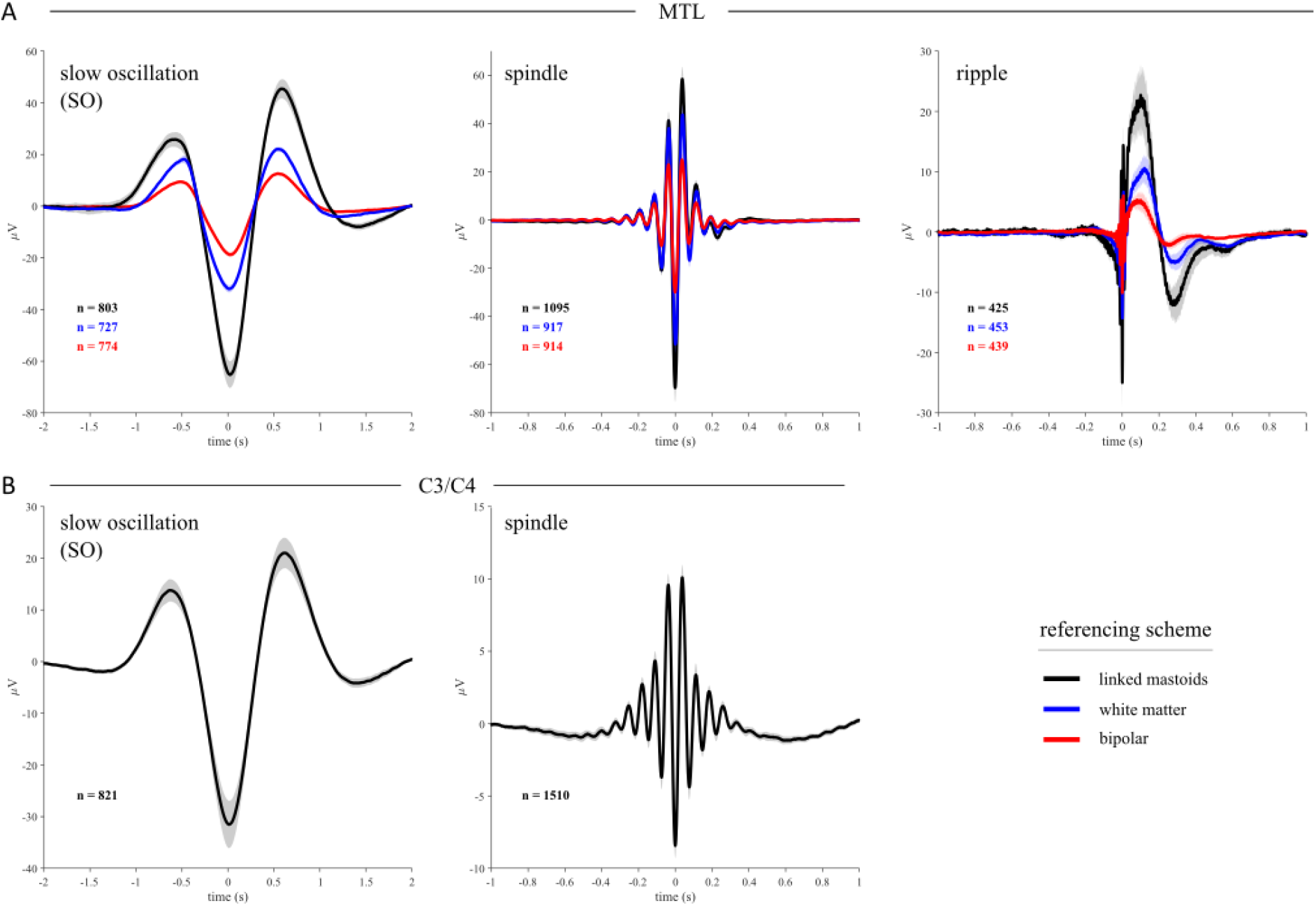
Effect of different referencing schemes on ERP amplitudes. Grand averages (n=20) ± SEM are shown, including the average numbers of detected events. **A**. Comparison of the effects of different (re-)referencing schemes on SOs, spindles and ripples in the MTL. *Black*: linked mastoid reference, *blue*: white matter contact on the same electrode, *red*: bipolar reference of adjacent distal electrode contacts. Note that despite stepwise amplitude decreases, the morphology and number of detected events remains similar. **B**. For comparison, SO and spindle grand averages ± SEM are shown collapsed across scalp electrodes C3 and C4 (referenced to linked mastoids only).

**Figure S2.**
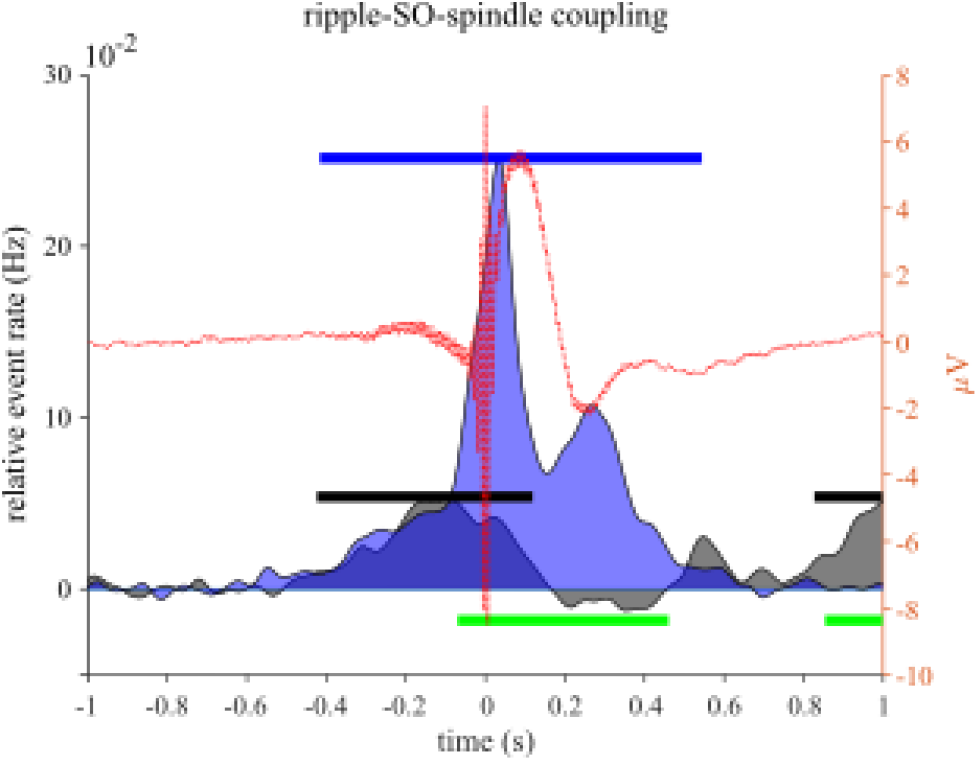
Spindle (blue) and SO (black) rates during ripples (dotted red line), relative to a pre-ripple baseline period. Note that unlike main Figure 2, event maximum times are plotted to better visualise SO up-states. Horizontal lines: blue … spindle occurrences vs. 0; black … SO occurrences vs. 0; green … spindle occurrences vs. SO occurrences, all P < .05 (corrected).

**Figure S3.**
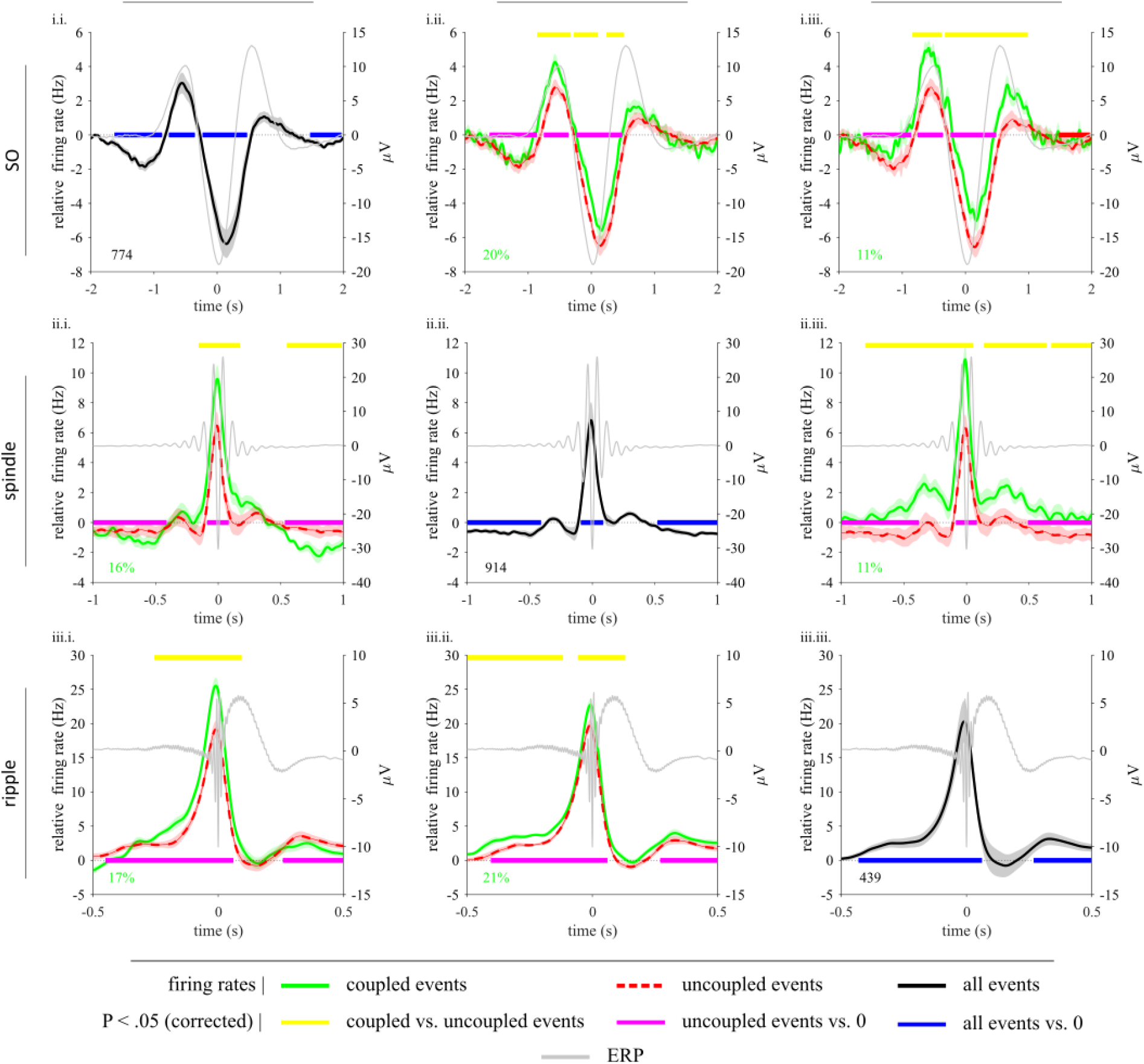
Event-locked neuronal firing rates (FRs) as a function of event contingencies (target event centre within ±1 s of seed event centre). *Rows*: seed events, *columns*: target events. *Diagonal*: all seed events (cf. main Figure 3A). Black lines represent mean baseline-corrected FR ± SEM of condition difference (coupled vs. uncoupled events). Grey lines represent grand average seed events (irrespective of target). Black numbers represent the number of events, averaged across contacts and sessions. Green numbers reflect percentage of coupled events (out of all seed events), averaged across contacts and sessions.

**Figure S4.**
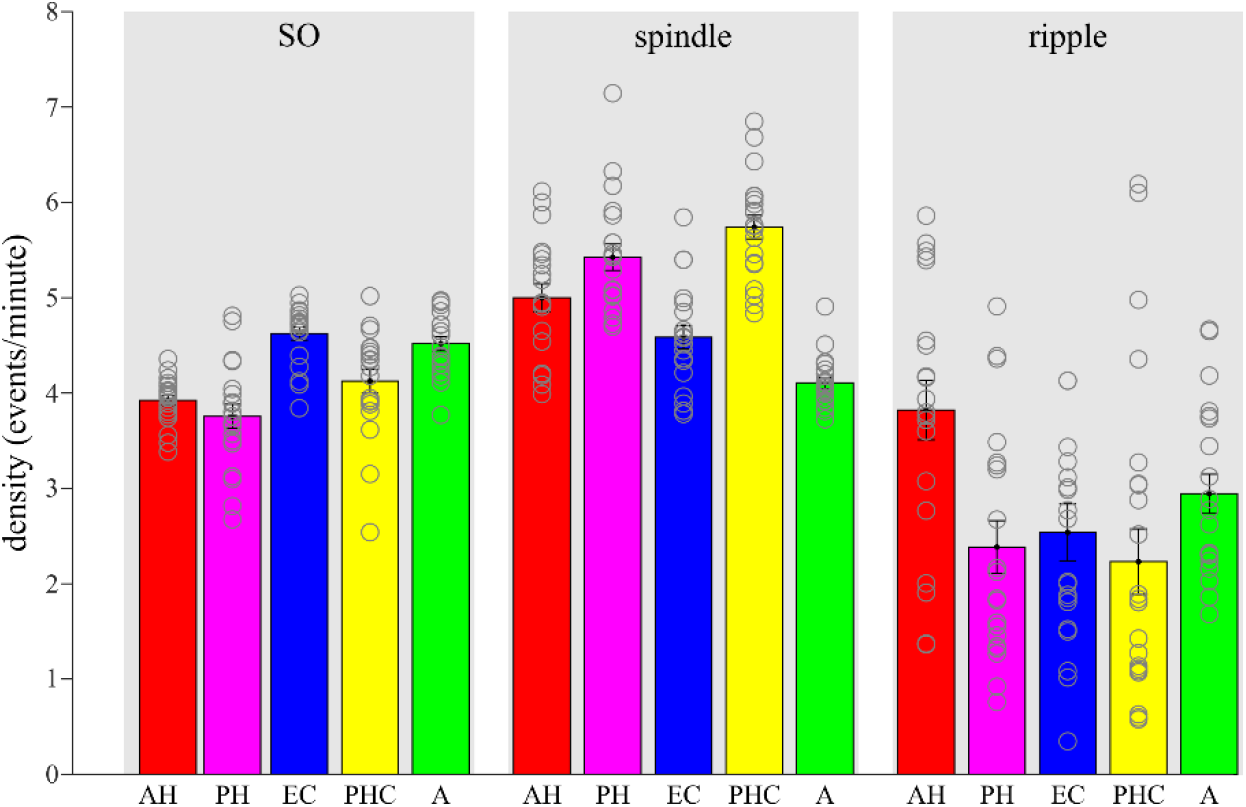
Event densities separated by MTL region. Bars represent mean ± SEM (n=20). Circles represent individual sessions. AH = anterior hippocampus, PH = posterior hippocampus, A = amygdala, EC = entorhinal cortex, PHC = parahippocampal cortex.

**Table S1.** Sleep architecture in minutes and percentages (mean and SEM across sessions, n=20). WASO … wake after sleep onset.

|  | minutes | % |
| --- | --- | --- |
| **N1** | 95 (8) | 18 (1) |
| **N2** | 174 (13) | 32 (2) |
| **N3** | 76 (5) | 15 (1) |
| **REM** | 75 (8) | 14 (1) |
| **WASO** | 114 (15) | 21 (3) |

